# Recurrent Neural Network-based Acute Concussion Classifier using Raw Resting State EEG Data

**DOI:** 10.1101/2020.07.07.192138

**Authors:** Karun Thanjavur, Arif Babul, Brandon Foran, Maya Bielecki, Adam Gilchrist, Dionissios T. Hristopulos, Leyla R. Brucar, Naznin Virji-Babul

## Abstract

Concussion is a global health concern. Despite its high prevalence, a sound understanding of the mechanisms underlying this type of diffuse brain injury remains elusive. It is, however, well established that concussions cause significant functional deficits; that children and youths are disproportionately affected and have longer recovery time than adults; and recovering individuals are more prone to suffer additional concussions, with each successive injury increasing the risk of long term neurological and mental health complications. Currently, concussion management faces two significant challenges: there are no objective, clinically accepted, brain-based approaches for determining (i) whether an athlete has suffered a concussion, and (ii) when the athlete has recovered. Diagnosis is based on clinical testing and self-reporting of symptoms and their severity. Self-reporting is highly subjective and symptoms only indirectly reflect the underlying brain injury. Here, we introduce a deep learning Long Short Term Memory (LSTM)-based recurrent neural network that is able to distinguish between healthy and acute post-concussed adolescent athletes using only a short (i.e. 90 seconds long) sample of resting state EEG data as input. The athletes were neither required to perform a specific task nor subjected to a stimulus during data collection, and the acquired EEG data was neither filtered, cleaned of artefacts, nor subjected to explicit feature extraction. The LSTM network was trained and tested on data from 27 male, adolescent athletes with sports related concussion, bench marked against 35 healthy, adolescent athletes. During rigorous testing, the classifier consistently identified concussions with an accuracy of >90% and its ensemble-median Area Under the Curve (AUC) corresponds to 0.971. This is the first instance of a high-performing classifier that relies only on easy-to-acquire resting state EEG data. It represents a key step towards the development of an easy-to-use, brain-based, automatic classification of concussion at an individual level.

## 1 Introduction

Concussion or mild traumatic brain injury (mTBI) is an urgent public health concern. Canadian [1] and US data [2, 3] indicate an annual rate of reported mTBIs of 1,100 per 100,000 people, 75% of which involve children, youths and young adults [4]. The actual rate is likely much higher as individuals suffering from concussion often do not seek medical advice [5, 6]. Despite its high prevalence, a sound understanding of the mechanisms underlying mTBI remains elusive. What is clear is that concussions induce diffuse, heterogeneous, spatially distributed changes in brain structure and function, resulting in cognitive, motor, emotional and behavioral challenges that can persist for many months [7–9]. Children and youths are especially vulnerable: They are disproportionately affected by concussions [10] and take longer than adults to recover [7,11]. The effects of concussion on their developing brain can lead to compromised brain and mental health, impairing learning, working and socializing [12]. Such disruptions at this critical developmental period in the lives of children and youths can be devastating.

The management of concussion suffers from a key challenge: There is no accepted, objective, brain-based tool to make an initial diagnosis of concussion. The current gold standard involves a clinical assessment by an experienced physician. This approach is difficult to standardize as it is dependent on the individual physician and relies on subjective reporting of the symptoms. Symptoms, however, do not appear to be directly correlated to the patho-physiological mechanisms responsible for the cognitive, emotional and behavioral deficits [13, 14] and several studies [14–23] have shown that structural and functional abnormalities in the brain persist even after symptom resolution, especially in pediatric patients. Manning et al. [20] and Hristopulos et al. [24] hypothesize that the increased vulnerability of children and youths to brain injury, as well as their longer recovery time, are consequences of the effects of concussion being overlaid on brains whose structural and functional organization are undergoing dynamic changes due to development. Moreover, recovering individuals are also at a higher risk of sustaining additional concussions, which can lead to neuro-psychiatric and neuro-degenerative disorders [25, 26]. Consequently, clinicians often find themselves in a quandary when trying to decide when pediatric patients can safely return to play/learn.

Numerous groups have sought to leverage neuroimaging data to not only detect mTBIs reliably and objectively, but also track brain recovery. At the group-level, structural changes in the integrity of white matter in adolescents has been observed using diffusion tensor imaging (DTI) [20, 27–32]. Similarly, studies of youths during “resting state” reveal significant mTBI-induced alterations in the functional organization of the brain with respect to their healthy counterparts, with key features being (i) a shift in spectral profile toward higher frequencies in the frontal brain regions, [33] (ii) an increase in functional connectivity (hyperconnectivity) [17, 34, 35], and (iii) disrupted information flow patterns [24], again particularly in the frontal regions. All of these studies rely on summary measures to characterize functional changes in the brain. However, because the brain injury due to mTBI is diffuse and the resulting changes in the brain structure and function are subtle, the summary measures cannot be used to statistically differentiate between healthy and injured at the *individual* level, leading the Radiological Society of North America to advise that “in pediatric patients, imaging should not be routinely obtained to diagnose mild TBI” because “there is insufficient evidence [to support] the routine clinical use of advanced neuroimaging techniques for diagnosis and/or prognostication at the individual patient level” [36].

Rapid advances in machine learning (ML) and artificial intelligence (AI) methods have opened up a new way to explore neuroimaging data [37–39]. ML evolved from the study of computer vision and computational learning theory, and is a method of data analysis that automates pattern recognition and model building. ML systems learn to identify relevant, often latent, features directly from the data, rendering explicit programming unnecessary. Consequently, they are especially suited for analysis of complex, high-dimensional datasets that themselves are products of often poorly understood dynamics. Focusing specifically on mTBI, several groups have attempted to develop ML algorithms to detect concussions. Examples of such attempts include the use of support vector machine (SVM) to distinguish between healthy and concussed individuals using explicit features derived from resting state magnetoencephalography (MEG) data [40] or resting state electroencephalography (EEG) recordings [41]; the use of random forest [42] and SVM [43] to classify subjects on the basis of structural connectivity features derived from diffusion magnetic resonance imaging (dMRI) data, with Vergara et al. [43] augmenting the dMRI measures with functional magnetic resonance imaging (fMRI)-based resting state functional network connectivity measures; the use of SVM to classify young adults using features from multiple task-related EEG recordings [44]; the use of an ensemble of shallow multilayer perceptron network with one hidden layer to distinguish between healthy and concussed adolescents using descriptors derived from resting state EEG data [45]; and most recently, the use of a deep learning convolution neural network to classify adolescents using EEG recordings of single-trial event-related potentials [46].

The goal of our research is to develop a reliable, accurate, easy-to-use deep learning neural network-based classifier (hereafter, *Conc*ussion *C*lassification *Net*work or *ConCNet*) that can distinguish between healthy and acute post-concussed individuals using only a *short* (i.e. 90 seconds long) sample of *raw resting state EEG* data (hereafter, raw90-rsEEG) as input. Here, “raw” refers to data that has neither been filtered nor cleaned to remove known artifacts; we discuss this further below. We explain our decision to use 90-second long samples in §4. Our decision to select EEG as the modality of choice was informed by a number of considerations, including the fact that (i) it is an inexpensive, non-invasive, direct, fine time resolution probe of the neural activity in individuals; (ii) at the group-level, it can reliably and objectively detect functional changes in the brain due to mTBI [24, 28, 47, 48]; and (iii) it does not require elaborate infrastructure, the introduction of contrast agents and radioisotopes, or exposure to radiation. Additionally, resting state data is easy to collect; the participant is neither required to perform a specific task nor subjected to a stimulus.

One challenge of working with EEG time series, particularly resting state EEG signals, is that they do not consist only of electrical signals from the brain, but also a variety of contaminants and distortions from environmental (e.g. strong power line interference at 60 Hz) and physiological sources (e.g. electrical signals generated by eye and head movements, eye blinks and cardiac activity). Cleaning the data involves identifying and removing these without altering the signals from the brain. Strategies in use range from fully automated algorithms to mixed schemes where some artifacts are removed via automated algorithms and others are identified and removed manually. The lack of a standardized, consensus cleaning pipeline and more importantly, the lack of well-defined measures for quantifying a dataset’s degree of “cleanliness” pose a serious challenge [49–51]. Additionally, commonly used artifact removal schemes are designed to treat known distortions, leaving open the possibility that the data may still be contaminated by latent artifacts. Even in the case of known artifacts, the identification-decontamination strategies are based on specific assumptions about the morphology of the distortions and the manner in which they interact with the signals of interest. These do not always fully capture the diverse ways in which the artifacts are manifest in real recordings [50, 52], and improper cleaning can introduce new distortions and unknown biases [50, 53]. Meisler et al. [54] assert that commonly used data cleaning methods do not necessarily improve brain state classification. In deliberately choosing to work with *raw* data, we circumvent these concerns and instead rely on the deep learning network to learn the relevant aspects of the data while ignoring the rest. In the process, we also avoid having to grapple with the complex problem of feature selection [55–57], which itself can be a source of bias [58–60], as well as confounding effects due to the EEG signals’ strongly non-stationary character on the feature values [61, 62].

In designing our classifier, we have adopted a recurrent neural network architecture comprising two layers of bi-directional Long Short Term Memory (LSTM) [63, 64] units. An LSTM-based network is ideally suited for processing an ordered sequence of data points, such as the EEG time series data, where there is a high likelihood of temporal correlations and/or causal relationships between measurements at different time points and where the time lag between these could potentially be large [65]. For a more detailed exposition of the reasons informing our decision to adopt a bi-directional LSTM network, including recent findings from the studies of brain-computer interface systems, we refer the reader to §4. As for training and testing the network, we used a total of 216 EEG samples obtained from 27 male adolescent athletes with sports related concussion within 1 month of injury, who met the concussion diagnostic criteria consistent with the Berlin consensus statement [66], and 280 datasets from 35 age and sex matched healthy athletes. For the concussed participants, the date and time of the direct blow was documented by the team coach as per the consensus statement, and the team physician or an experienced physician with expertise in concussions made the diagnosis of concussion based on the Berlin consensus statement. The number of symptoms and symptom severity of each concussed subject were assessed using either the Sports Concussion Assessment Tool 3 (SCAT3) or the Child Sports Concussion Assessment Tool 3 (Child SCAT3) (see §4).

## 2 Results

In Table 1, we present the demographic information about the participants in this study. All of the concussed participants met the Berlin criteria and exhibited from 4 to 22 SCAT3 symptoms, at the time of testing. The symptoms most frequently reported included irritability, sensitivity to light, dizziness, fatigue, “don’t feel right”, and difficulty concentrating/remembering.

**Table 1.** Demographic information for the participants involved in this study. SD stands for standard deviation.

| Demographic information | Healthy | Concussed |
| --- | --- | --- |
| Age in years (SD) | 14.7 (2.1) | 13.5 (2.6) |
| Sex | 100% male | 100% male |
| Time since concussion |  | 100% within one month |
| SCAT: Number of symptoms (SD) |  | 9.2 (6.2) |
| SCAT: Symptom severity (SD) |  | 21.7 (18.7) |
| Child SCAT: Number of symptoms (SD) |  | 13.0 (5.4) |
| Child SCAT: Symptom severity (SD) |  | 23.2 (13.6) |

The development and assessment of *ConcNet* proceeded in three stages. Here, we briefly summarize these three stages, focusing on the results. Additional details are provided in the Methods section (§4).

### Stage 1: Identifying a promising machine learning algorithm for *ConcNet*

A number of different machine learning techniques have been proposed to tackle classification problems (see Kotsiantis et al. [67] and Singh et al. [68] for comprehensive reviews). Prior to settling on a deep learning neural network-based classifier, and specifically an LSTM-based deep learning recurrent neural network, we experimented with other options, including using Support Vector Machine (SVM) to distinguish between healthy and concussed individuals on the basis of features derived from filtered and cleaned, resting state EEG data [41], and using an ensemble of multilayer perceptron networks to distinguish between the two populations on the basis of descriptors derived from filtered but not cleaned resting state data [45]. In all cases, the classifiers were trained and validated using a subset of the available data; tested on the remaining data using a *“blinded”, one-shot* protocol, as described in §4; and evaluated on the basis of *Accuracy* and *Consistency. Accuracy* (and most of the other commonly used assessment metrics) are defined in Table 2 while *Consistency* is defined below.

**Table 2.**
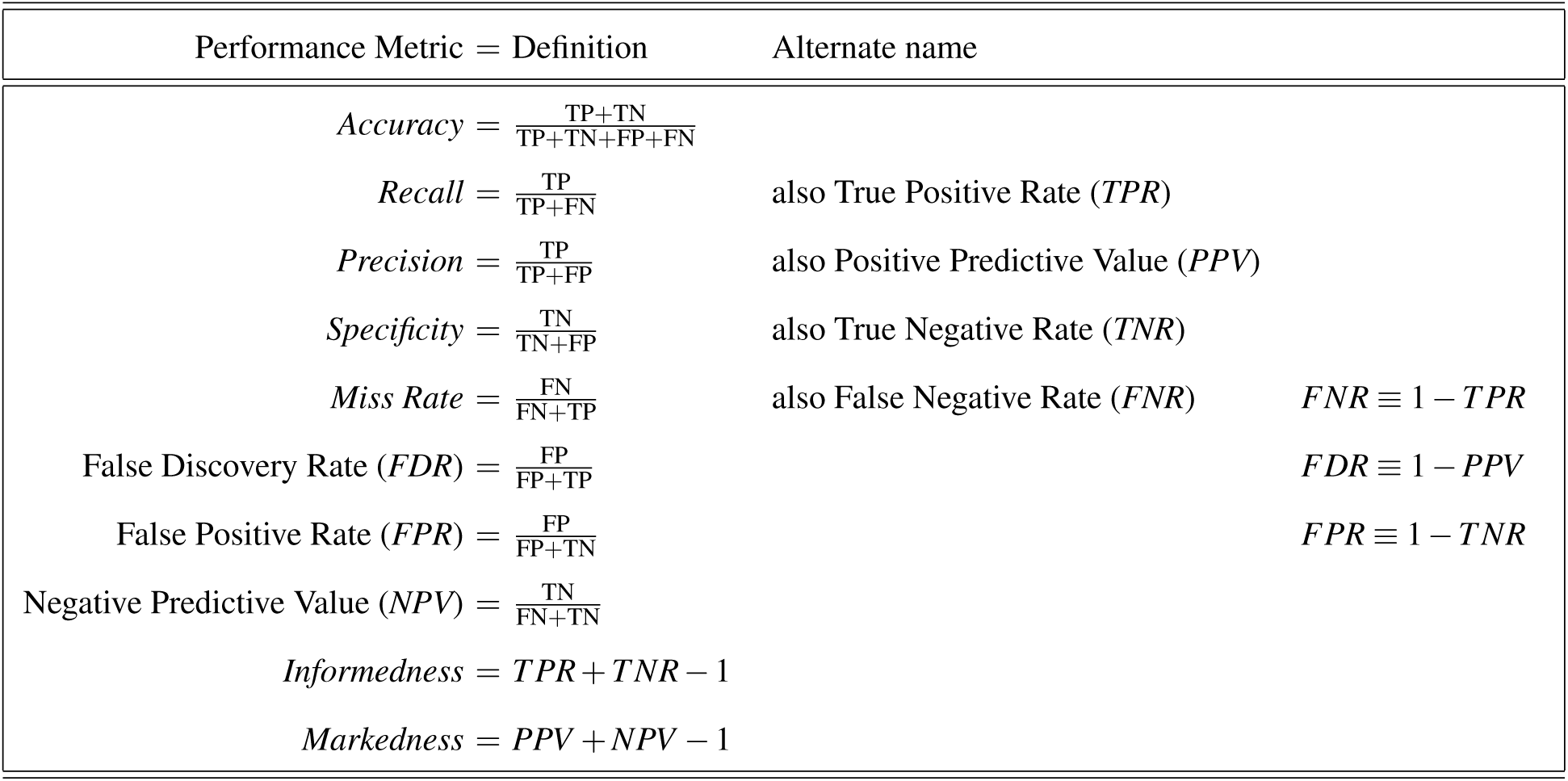
Common measures used in this paper to gauge the performance of classification tasks. In the formula above, TP (True Positives) is the number of correctly classified concussed individuals; TN (True Negatives) is the number of correctly classified healthy participants; while FP and FN are the numbers of falsely classified healthy (i.e. False Positives) and concussed (i.e. False Negatives) participants, respectively.

Since our aim is to develop a clinically viable classifier, we considered an algorithm as promising only if its *Accuracy* was > 80% during the Stage 1 testing phase. *Accuracy* is the fraction of classifications that are correct. All except the LSTM-based classifier failed to reach this threshold, with the features-based algorithms achieving an accuracy of at the most 65%. The LSTM classifier, the focus of this paper, achieved *Accuracy* = 88.9%. The architecture of our first generation *ConcNet*, following Stage 1 training and tuning, is shown in Figure 1.

**Figure 1.**
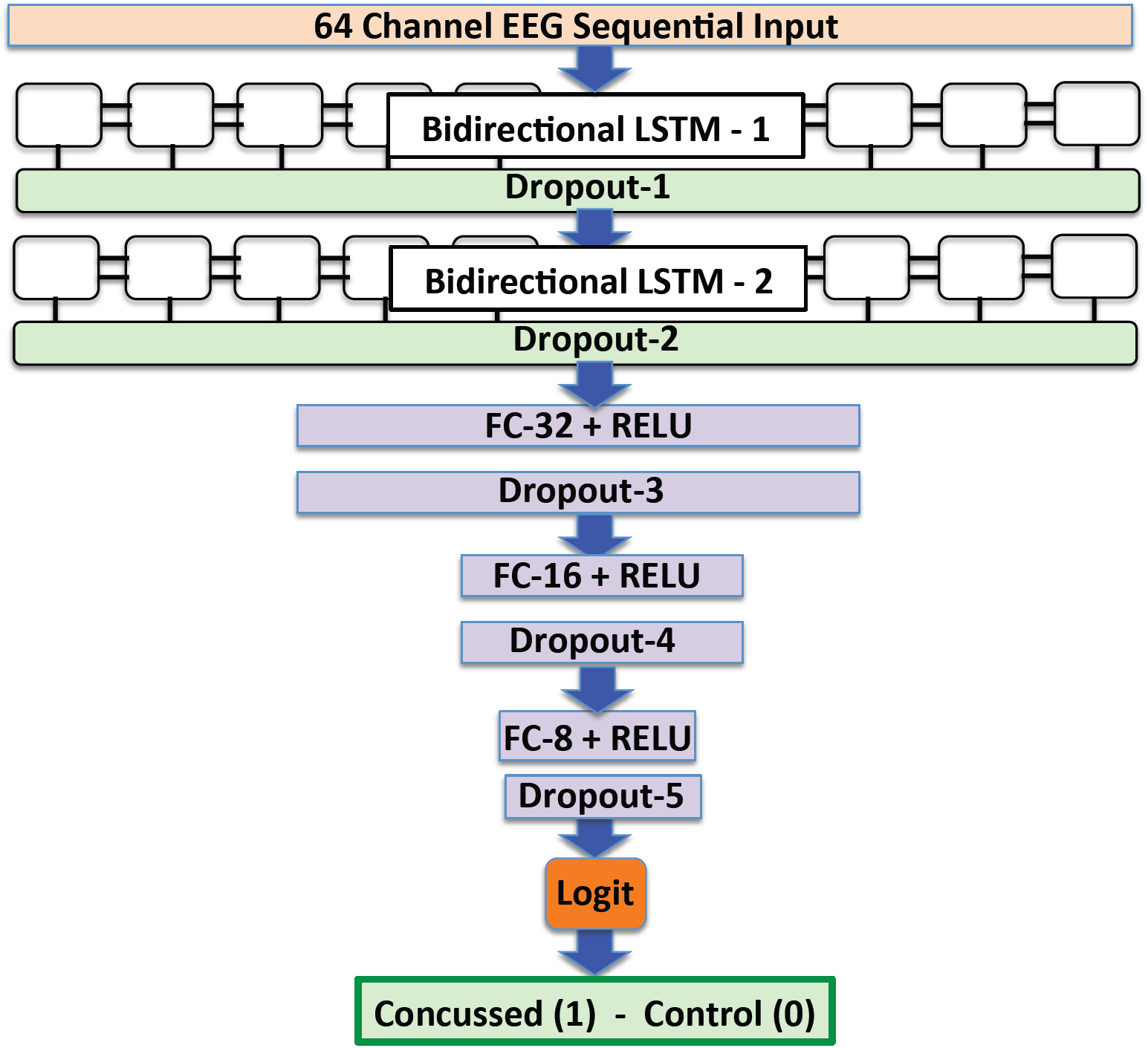
The architecture of our first generation *ConcNet*, a recurrent neural network architecture comprising two LSTM layers. The 64-channel EEG input is fed through a sequential input layer to the first layer with bi-directional Long Short Term Memory (LSTM) units. The output is then passed to a dropout layer for regularization to reduce overfitting, and then to the second LSTM layer that is similar to the first, another dropout layer, followed by three sets of fully connected and dropout layers. The fully connected layers have 32, 16 and 8 units and use a Rectified Linear Unit activation functions. The output is passed to a *softmax* logistic regression (logit) classifier, where it is assigned a probability and classified as *concussed* or *healthy*.

In *ConcNet*, each input is assigned a probability (*P*_mTBI_) by the *softmax* logistic regression (logit) unit at the output layer and by comparing this probability to the user-chosen classification threshold value, given a score of 0 (*healthy*) or 1 (*concussed*). For *ConcNet*, we set the threshold at 0.5 and a participant with a probability score of greater than this threshold (i.e. *P*_mTBI_ > 0.5) is classified as concussed. *ConcNet*’s performance after two rounds of Stage 1 blind test is shown in Table 3. The confusion matrix summarizing the results is shown in Figure 2. Apart from *Accuracy*, the model also achieved *Recall* = 100%, *Precision* = 80% and *Specificity* = 80%. *Recall* (also known as *Sensitivity* or True Positive Rate) measures the proportion of actual positives (i.e. concussed) that are correctly identified as such or alternatively, it is a measure of the classifier’s ability to identify concussed as concussed. *Precision* is the fraction of true positive in the set classified as positive (i.e. the set of both true and false positives); it is a measure of how much trust one can have in the classifier’s positive predictions. *Speci f icity* (also, True Negative Rate) measures the fraction of participants labelled as healthy that are correctly identified as such. The network correctly identified all of the concussed samples but misidentified three raw90-rsEEG samples contributed by three different healthy participants. In this regard, the classifier worked just as one would want a clinical classifier to, correctly identifying all the concussed subjects (i.e. high recall), and although a small number of healthy participants were misclassified (i.e. a slightly lower precision), it can be argued that this is an acceptable trade-off because *ConcNet* is erring on the side of caution.

**Table 3.**
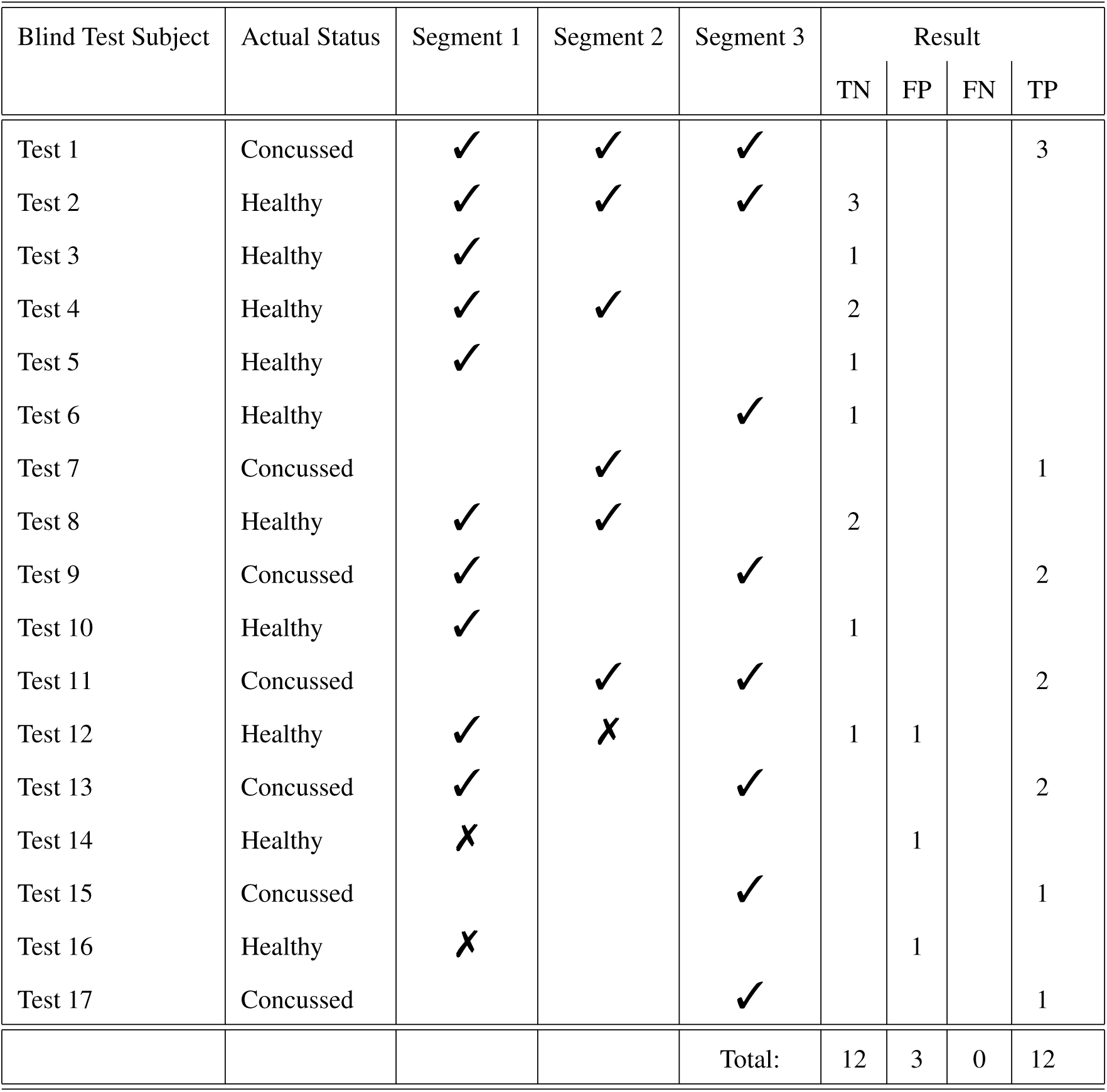
Details of the Stage 1 blind test results for *ConcNet*. The table shows the number of distinct segments from any one participant were that were present in the blind test sample, and how each of these were classified. All except one pair were consistently — and correctly — classified. Final column gives the results in terms of the number of True Negative (TN), False Positive (FP), False Negative (FN) and True Positive (TP). These are further summarized in the Confusion Matrix in Figure 2.

| Blind Test Subject | Actual Status | Segment 1 | Segment 2 | Segment 3 | Result |  |  |  |
| --- | --- | --- | --- | --- | --- | --- | --- | --- |
|  |  |  |  |  | TN | FP | FN | TP |
| Test 1 | Concussed | ✓ | ✓ | ✓ |  |  |  | 3 |
| Test 2 | Healthy | ✓ | ✓ | ✓ | 3 |  |  |  |
| Test 3 | Healthy | ✓ |  |  | 1 |  |  |  |
| Test 4 | Healthy | ✓ | ✓ |  | 2 |  |  |  |
| Test 5 | Healthy | ✓ |  |  | 1 |  |  |  |
| Test 6 | Healthy |  |  | ✓ | 1 |  |  |  |
| Test 7 | Concussed |  | ✓ |  |  |  |  | 1 |
| Test 8 | Healthy | ✓ | ✓ |  | 2 |  |  |  |
| Test 9 | Concussed | ✓ |  | ✓ |  |  |  | 2 |
| Test 10 | Healthy | ✓ |  |  | 1 |  |  |  |
| Test 11 | Concussed |  | ✓ | ✓ |  |  |  | 2 |
| Test 12 | Healthy | ✓ | ✗ |  | 1 | 1 |  |  |
| Test 13 | Concussed | ✓ |  | ✓ |  |  |  | 2 |
| Test 14 | Healthy | ✗ |  |  |  | 1 |  |  |
| Test 15 | Concussed |  |  | ✓ |  |  |  | 1 |
| Test 16 | Healthy | ✗ |  |  |  | 1 |  |  |
| Test 17 | Concussed |  |  | ✓ |  |  |  | 1 |
|  |  |  |  | Total: | 12 | 3 | 0 | 12 |

**Figure 2.**
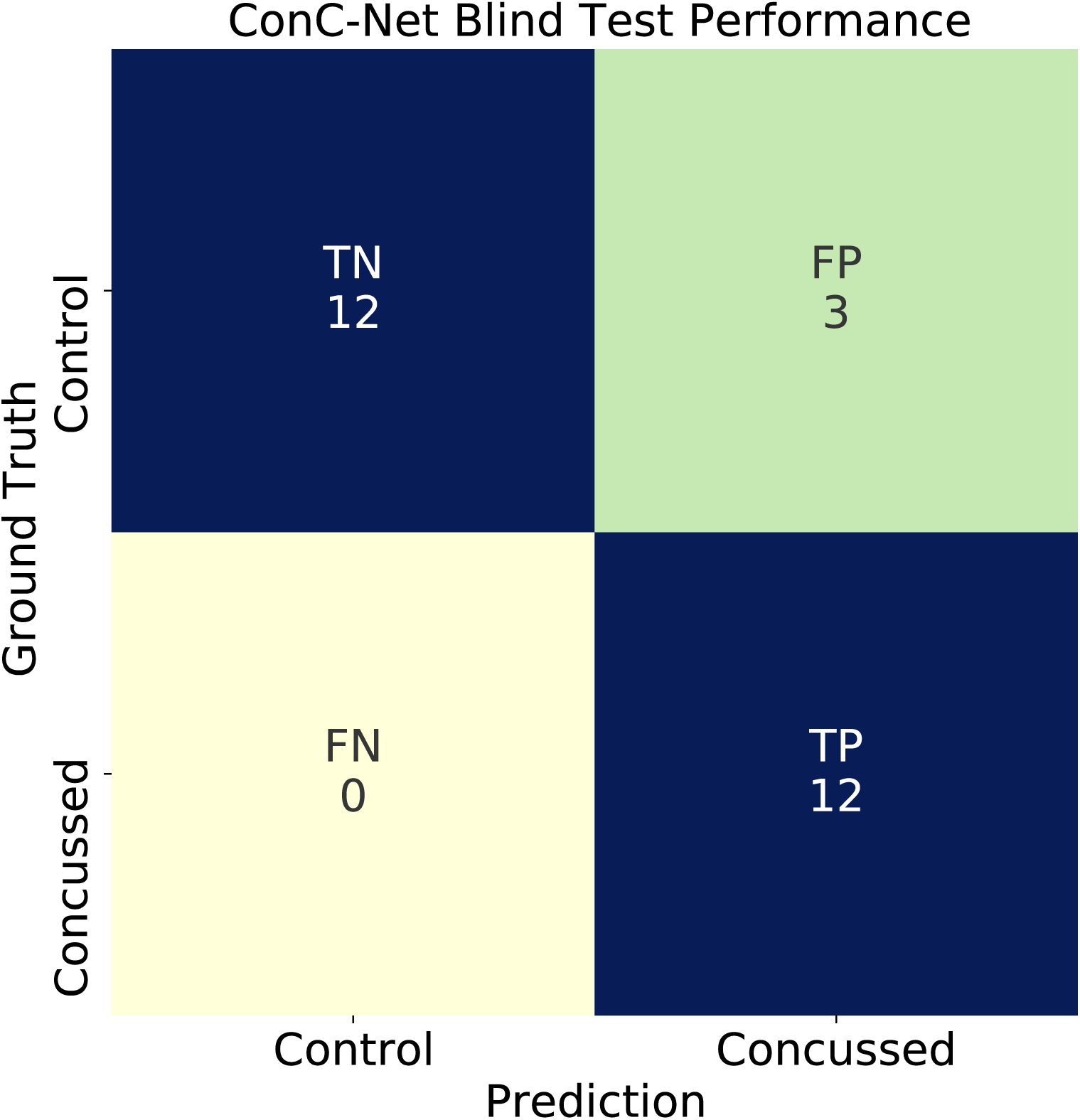
Confusion Matrix summarizing the results from the blind test carried out in Stage 1.

As illustrated in Table 3, the blind sample comprised more than one (but not identical) raw90-rsEEG samples from some of the subjects. Each sample consists of a segment of synchronous 90 second-long EEG time series. The motivation for including different segments taken from the same subject is to investigate if these are similarly classified (consistency). The classifier’s *Consistency* is defined as the ratio of the distinct number of segment pairs contributed by the same subjects that are similarly classified, to the total number of such pairs in the sample. Our blind sample included a total of 12 such pairs of raw90-rsEEG recordings, each pair derived from the same subject. The LSTM network classified 11 segment pairs identically (and correctly) for a *Consistency* rating of 91.7%.

### Stage 2: Optimizing the LSTM-based *ConcNet*

Encouraged by the Stage 1 results, we proceeded to systematically optimize and simplify *ConcNet*’s structure without compromising on performance. The final network architecture, shown in Figure 3, is largely the same as *ConcNet* 1.0 with one key difference. It has one less fully connected layer. Specifically, *ConcNet* 2.0 is made up of (1) a sequential input layer that receives the 64-channel raw90-rsEEG data and passes it to (2) the first LSTM layer comprising 100 memory units. The output is handed off to (3) a dropout layer that randomly discards the results from 30 of the 100 LSTM units (i.e. *DOf* =30%). Dropout is a commonly used regularization technique to prevent overfitting. The output from the Dropout layer is passed to (4) a second LSTM layer with 100 bi-directional memory units, then to (5) another dropout layer with *DO f* = 30%. The series data thus synthesized by the two LSTM+Dropout layers is fed to two fully connected (FC) layers, (6) the first with 8 nodes and (7) the second with 2 nodes, with each of the nodes utilizing the ReLU (see §4) activation function. Given the small number of nodes in these FC layers, the corresponding dropout layers present in *ConcNet* 1.0 (c.f. Figure 1) were no longer necessary. As noted previously, the binary classification and the assignment of a probability happens in the final *softmax* logit layer.

**Figure 3.**
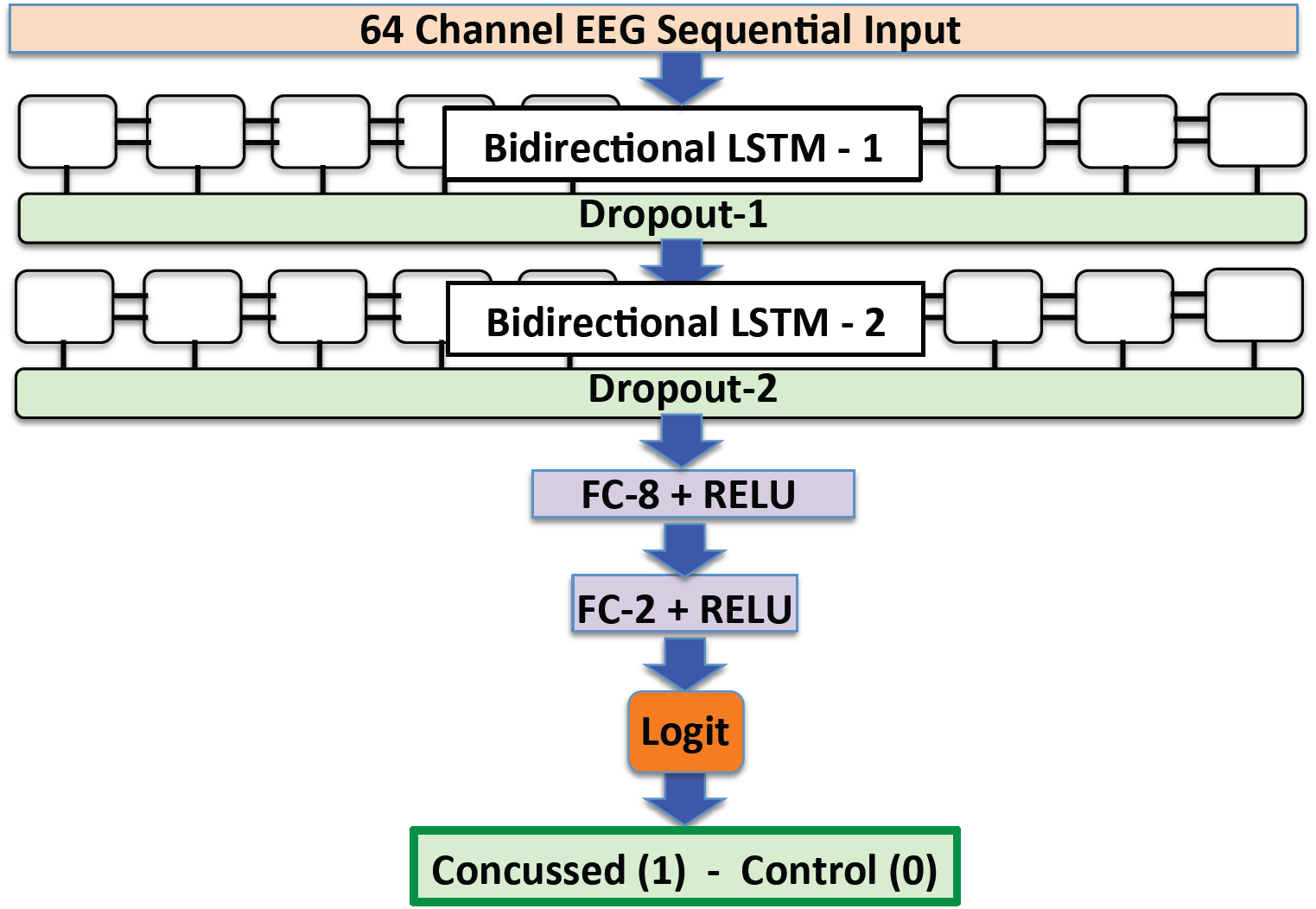
The architecture of our final streamlined version of *ConcNet* (*ConcNet* 2.0). The architecture is similar to that shown in Figure 1 except that the version shown here only 2 fully connected layers, with 8 and 2 units, and no associated dropout layers.

In addition to being somewhat more streamlined than our first generation version, our final network also formally fared a bit better in that the number of false positives dropped from 3 to 2 and correspondingly, the accuracy increased to 92.6%. We emphasize that this formal improvement in the performance is not statistically significant but it did mean that we had accomplished our stated goal: to streamline the network architecture as much as possible while maintaining performance. Going forward, the network architecture and hyperparameters were held fixed for the duration of the experiment.

### Stage 3: Assessing *ConcNet* ‘s Performance

The third and the final stage involved carefully assessing the performance of *ConcNet* 2.0, including quantifying the uncertainties associated with its performance metrics. We were specifically interested in assessing how well *ConcNet* would generalize as the training, validation and testing datasets are varied. To that end, we drew upon the full dataset of available raw90-rsEEG recordings and used stratified balanced sampling strategy (see §4 for details) to train 100 independent *ConcNet* 2.0 networks; that is, networks that are identical copies, in terms of architecture and hyperparameters, of the final network configuration from Stage 2 above.

Table 4 shows the median values (*Q*_2_), as well as the 25% (*Q*_1_) and 75% (*Q*_3_) quartiles, of the metrics commonly used to gauge a classifier’s performance. Notably, the *Accuracy* and *Speci f icity* remain high, with median values at 0.917 (*Q*_1_ = 0.833, *Q*_3_ = 1.000) and 0.900 respectively (*Q*_1_ = 0.850, *Q*_3_ = 1.000) respectively. All three *Recall* quartiles are equal to 1, while the mean is also reassuringly high at 0.93.

**Table 4.** The ensemble median (*Q*_2_), the 25% percentile (*Q*_1_), and the 75% percentile (*Q*_3_), of the various network metrics distributions. The statistics are derived from the 100 networks ensemble test, quantifying the performance of *ConcNet* 2.0. The results for the five participants who were systematically misclassified have *not* been excluded.

| Metric | $Q_2$ | $Q_1$ | $Q_3$ | Metric | $Q_2$ | $Q_1$ | $Q_3$ |
| --- | --- | --- | --- | --- | --- | --- | --- |
| *Accuracy* | 0.917 | 0.833 | 1.000 | *Area Under Curve (AUC)* | 0.971 | 0.963 | 0.975 |
| *Recall (TPR)* | 1.000 | 1.000 | 1.000 | *Informedness* | 0.900 | .800 | 1.000 |
| *Precision (PPV)* | 0.667 | 0.500 | 1.000 | *Markedness* | 0.667 | 0.500 | 1.000 |
| *Specificity (TNR)* | 0.900 | 0.850 | 1.000 | *Miss Rate (FNR)* | 0.000 | 0.000 | 0.000 |
| *False Discovery Rate (FDR)* | 0.333 | 0.000 | 0.500 | *False Positive Rate (FPR)* | 0.100 | 0.000 | 0.150 |

We also examined the Receiver Operating Characteristic (ROC) curve and the corresponding Area Under the Curve (AUC) for *ConcNet*. The latter is a particularly useful measure because while the values of *Accuracy, Specificity* and *Recall* depend on the adopted threshold for classification, AUC does not. An ROC curve plots the True Positive Rate (TPR) vs. the False Positive Rate (FPR) for a range of classification thresholds. In the present case, we vary the probability threshold for being classified as concussed from 0-100%. A low classification threshold generally leads to more healthy participants being classified as concussed. Figure 4 shows the ROC derived using the results of the 100 networks. The ensemble median AUC for *ConcNet* is 0.971 (*Q*_1_ = 0.963, *Q*_3_ = 0.975). AUC measures the performance across all classification thresholds: An AUC of 0.5 corresponds to a classifier whose accuracy is no better than a coin toss while an AUC of 1.0 corresponds to a perfect test (i.e. 100% recall and 100% specificity). A value of > 0.96 indicates a classifier with excellent discriminatory ability [69].

**Figure 4.**
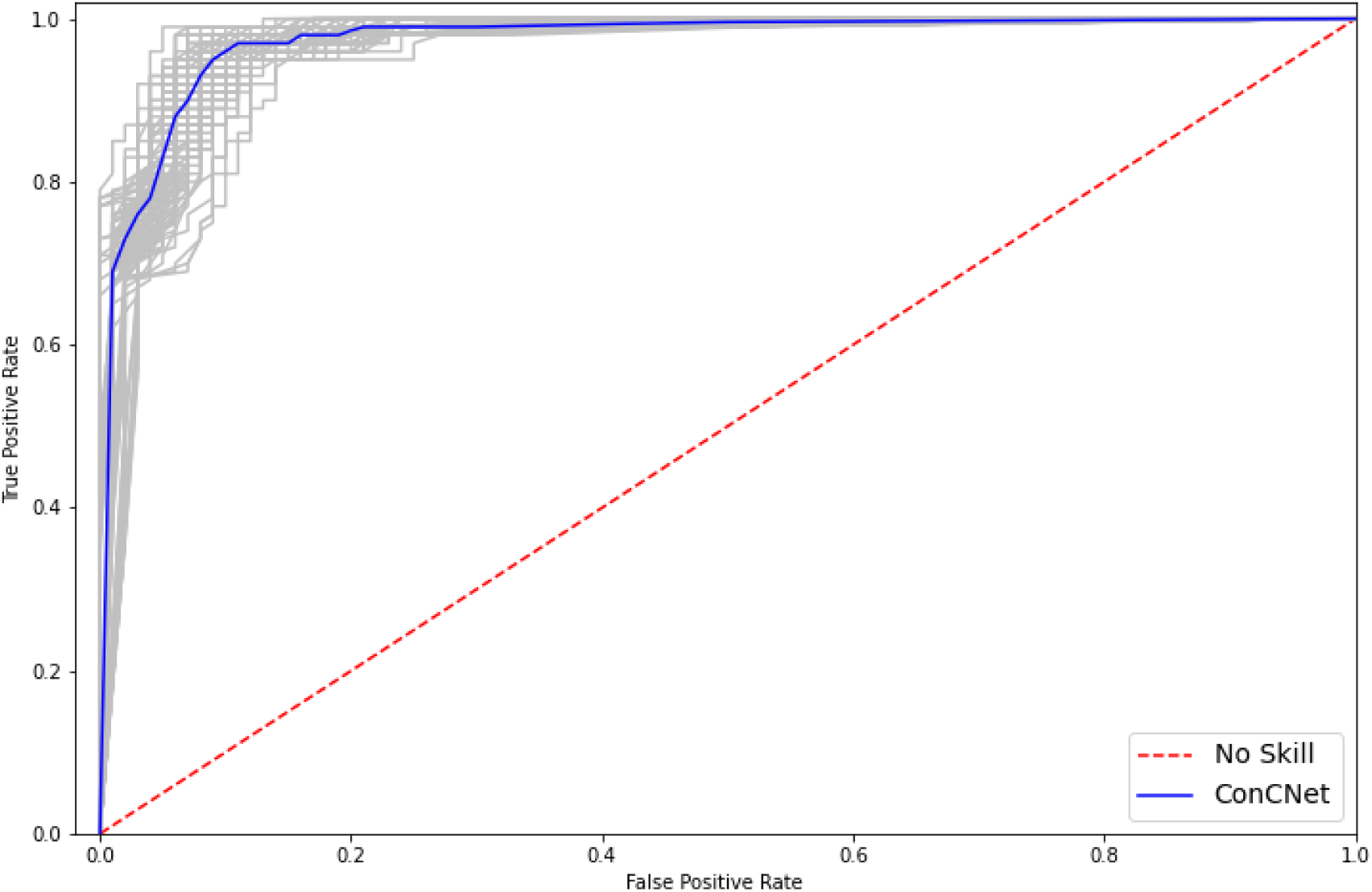
The Receiver Operating Characteristic (ROC) curves for the 100 LSTM networks. The grey curves show the results for each of the 100 networks and the blue curve shows the median. The ensemble median Area Under the Curve (AUC) is 0.971 (the 25% and the 75% percentile are 0.963 and 0.975, respectively). A classifier with no better accuracy than chance would be expected to have an AUC of 0.5, while an AUC of 1 corresponds to a classifier with perfect accuracy. A value of > 0.96 indicates a classifier with excellent discriminatory ability [69].

To investigate *ConcNet* results further, we examined the full set of predicted scores assigned to each subject. We note that a subject is assigned a score only when a raw90-rsEEG recording from that subject is presented to a network as a test case. Figure 5 shows the median of the predicted scores as well as the corresponding 25^*th*^ and 75^*th*^ percentiles. For clarity, the results are divided across two panels, with the left panel showing the results for the 27 concussed participants and the right panel summarizing the results for the 35 healthy participants. In each panel, the blue points represent those subjects who were correctly classified. The total number of points may seem to be fewer than expected, given the number of participants in each category; however, this is not the case. Many of the blue points in the *true positive* (TP) and *true negative* (TN) quadrants have very similar scores and are stacked on top of each other. The small error bars associated with these points indicate that the corresponding subjects were assigned similar predicted scores regardless of which network from the ensemble was used or which of their eight raw90-rsEEG recordings was analyzed.

**Figure 5.**
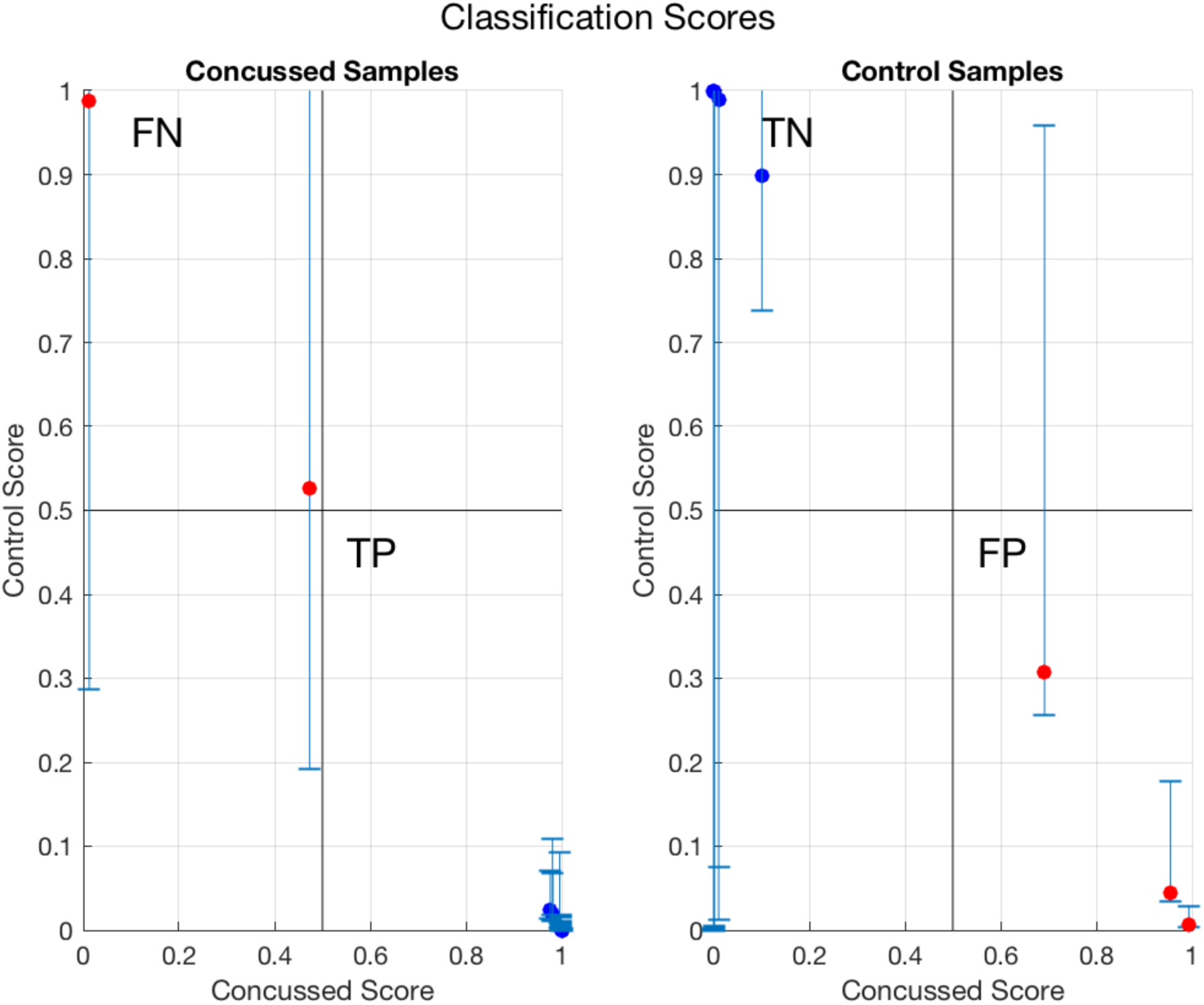
The median value of the probabilities, *P*_mTBI_, assigned by the *softmax* logistic regression (logit) unit to each concussed and healthy participant whenever the sample was used for testing a network in the ensemble, plotted along the *x* − axis, and the median value of *P*_Hlth_ ≡ (1 − *P*_mTBI_) plotted on the *y* − axis. The − *y* error bars show the 25^*th*^ and 75^*th*^ percentiles of the predicted probability scores. The left panel shows the results for the 27 concussed subjects, while the right shows the results for the 35 healthy individuals. Blue points represent samples which were properly classified based on our criterion of *P*_mTBI_ > 0.5. The red points correspond to the individuals whose median score resulted in a misclassification.

As for the misclassifications, of the 27 concussed subjects, we found that one was nearly always (i.e. consistently) misclassified while the EEG samples of another subject tended to be misclassified more often than not (and therefore, the median fell below our threshold). These two cases appear as red points in the false negative (FN) quadrant of the left panel in Figure 5. As for the healthy subjects, we found that two were misclassified most of the time and one was more frequently misclassified than not. These are shown as red points in the right panel of Figure 5. The scatter in the predicted scores for all of the misclassified cases is generally large.

Analyzing the misclassifications further, we noted that they were *not random occurrences* in the sense that the same five individuals tended to be misclassified regardless of which of the subjects’ raw90-rsEEG recordings was used or the network in the ensemble it was processed through. To further illustrate this, in Figure 6, we show the scores as stacked bar charts for all the concussed and healthy samples. For each bar, the blue and red areas represent the fractional number of times that the subject under consideration was respectively correctly or incorrectly classified. Any bar whose red portion occupies >50% of the column corresponds to a sample which was more frequently misclassified. The two concussed and the three healthy “problematic” subjects readily stand out.

**Figure 6.**
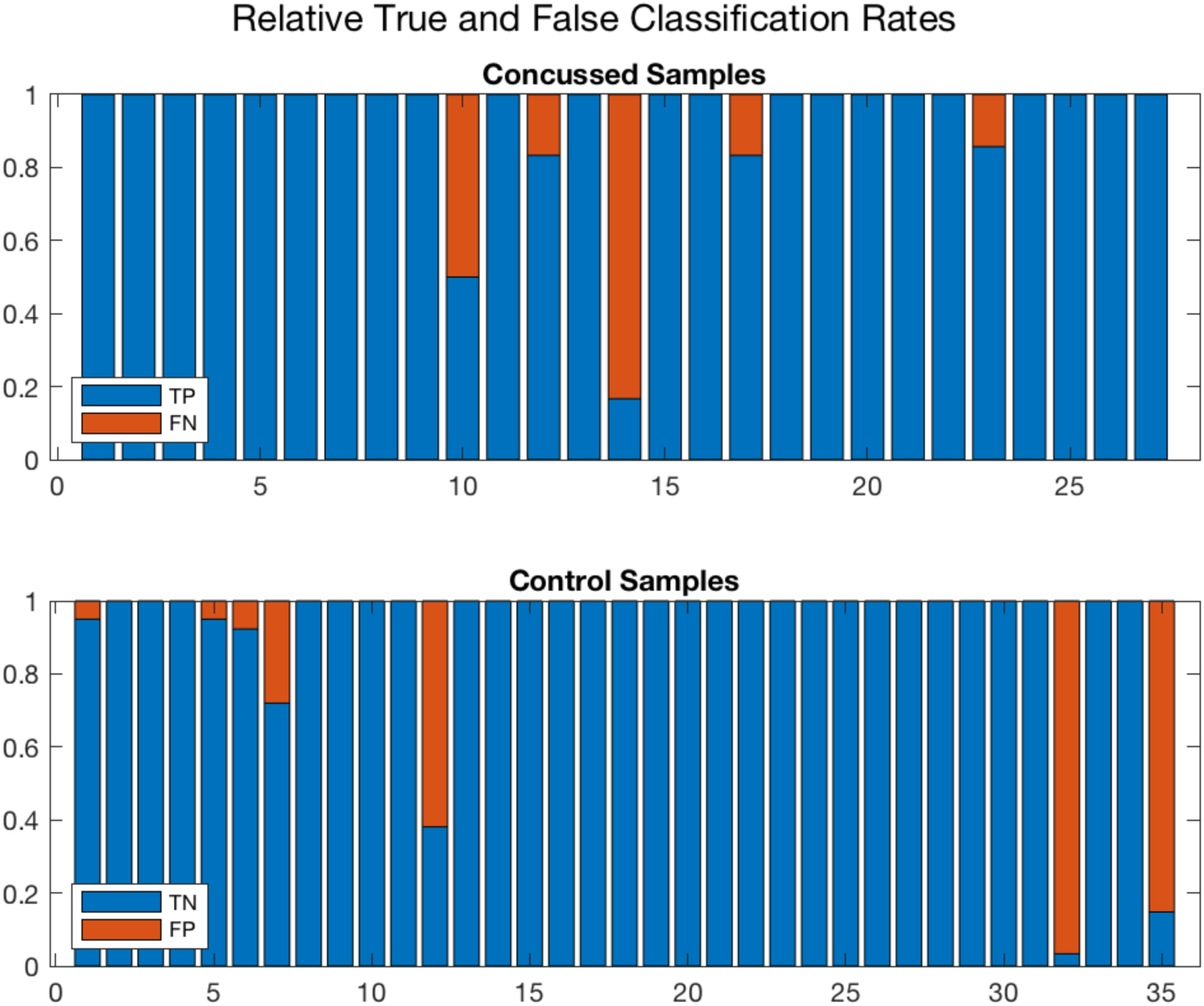
A different representation of the ensemble results for the 27 concussed and 35 healthy samples. The blue and red regions in each column illustrate the relative fractions of times that participant was properly or wrongly classified by the ensemble of networks. Three healthy and two concussed “problematic” subjects tended to be systematically misclassified by the networks in the ensemble. A handful of other participants were sometimes misclassified by one or two of the networks. On further examination, the training of the latter involved an EEG sample from one of the “problematic” individuals.

Figure 6 also shows a few additional columns with small red segments. These segments represent a small number of networks that misclassified the subjects in question. We found that these failures were also largely due to the ‘problematic” samples highlighted above. Since the partitioning of the full dataset into training, validation and testing for each of the 100 networks is done at random, every so often the EEG data from one or more “problematic” cases appeared in the training/validation subsets for some of the networks. This degraded the performance of the affected networks. The moderate value for the *Precision* and *Markedness*, and the spread in their values, are primarily due to the five “problematic” individuals.

The systematic nature of the problem makes us suspect the issue is not with the classifier but with the data. We will return to this matter in §3 and discuss possible reasons for the misclassifications. As a test, we removed the five systematically misclassified individuals from our dataset, and retrained and tested a single network. Not surprisingly, the overall accuracy rose to 98%, an improvement of nearly 5% from our previous Stage 2 results.

## 3 Discussion

One of major current gaps in concussion management is determining when an athlete has recovered from the initial injury after a sports related concussion. Health care providers often have to make difficult decisions regarding when an athlete is fit to safely return to sport/activity based on self-reporting of symptoms and their severity, and clinical testing. Self-reports, however, are highly subjective and also influenced by socio-cultural factors [70, 71]. Moreover, emerging literature is showing that neurophysiological changes persist well after clinical recovery [15–23], suggesting that athletes may be returning to activity before their brains have fully recovered and may therefore be at increased risk of additional brain injury [66]. In fact, a history of prior concussions is one of the strongest predictors of future concussions [71–73]. Specifically, high school athletes with prior concussions have >2 × higher risk of suffering subsequent concussions [72, 73] and with each successive injury, have a growing risk for sustaining long term neurological and mental health complications. Brain based measures of neurophysiological recovery (based on fMRI, DTI and EEG) have so far only been able to detect differences in the brain at a group level. The results presented here demonstrate that *ConcNet*, our LSTM-based deep learning network for concussion classification, is a promising engine for an objective, brain-based platform for identifying acute post-concussed individuals using only their resting state EEG data. Admittedly, the EEG data used to train and test the network was acquired from only male adolescent athletes (ages: 10–18), and this is a limitation of the present study. Nonetheless, *ConcNet*’s excellent performance serves as a proof of concept demonstrating that it is possible to train a deep learning network to recognize acute post-concussed individuals, with a high degree of accuracy, using only their resting state EEG data.

Encouraged by our current results, we were actively expanding our database to include female athletes as well as younger and older subjects until interrupted by the Covid-19 pandemic. Several studies find that female athletes have an inherent elevated risk for concussion compared to their male counterparts, differ in the types (and severity) of symptoms experienced, and take longer to become symptom free [70, 74–76]. However, in the absence of an objective tool for confirming a concussion, it is difficult to establish whether this is a genuine difference or the result of a bias related to how males and females report concussions and concussion symptoms [70, 74]. *ConcNet*-like platform offers an opportunity to expand the current understanding of how sex, among other factors, affects injury epidemiology and outcomes.

Of the 27 concussed subjects considered in this study, 25 subjects were consistently classified correctly while 2 subjects were tended to be misclassified as healthy. We have already noted that adjusting the classification threshold slightly would have reduced the number of misclassifications to 1; however, at the moment, we do not have a clear basis for doing so. Moreover, we are intrigued by the systematic differences in the probabilistic scores assigned to the 2 individuals in questions, compared to the scores assigned to the other 25 subjects. Three possible explanations stand out:

Firstly, we cannot outrightly exclude the possibility that the failures reflect *ConcNet*’s small but finite probability of misclassifying. However, since the misclassifications are not randomly distributed across the subjects and that most of the misclassifications are associated with the raw90-rsEEG datasets contributed by the five problematic subjects, we find it hard to accept that these are products of random errors.

Secondly, the two subjects may have been misdiagnosed at the outset. Our criteria for identifying subjects with concussion were (i) documentation of the date and time of a direct blow by the team coach, and (ii) a diagnosis by an experienced physician. An error would have had to have happened at both stages and while we cannot exclude the possibility, we think it is also unlikely.

The third possibility is that either the subjects were well on their way to recovering from their injury at the time their brains were scanned or that their brains were only minimally affected by the injury. In reviewing the two misclassified concussed subjects’ other available data, we found that they were indistinguishable from the other concussed subjects except for the fact that they were both scanned at 1 month post injury, and comprised 2 of the 3 individuals in our sample at the edge of our time window, which lends itself to the “on the way to recovery” conjecture. All the other subjects were scanned at 1 week post injury. In light of this, perhaps a more pertinent question is not why the two subjects were misclassified but rather, why was the third subject in the 1-month post injury cohort not misclassified. We speculate that the “correctly” classified member of the cohort may have suffered a more pronounced injury or may be experiencing a more prolonged recovery. Neuroimaging studies find that the brain’s response to an mTBI, as suggested by a significant spread in the measures derived from neuroimaging probes of the brain’s structure and function, varies considerably from subject to subject both at the acute phase as well as in the brain’s recovery trajectory [77, 78]. In fact, this same explanation may also explain the three subjects in the healthy group that were misclassified as concussed. Longitudinal neuroimaging studies show that in some individuals, mTBI induced changes in the brain can persist for several months and even past a year [15–23]. These persistent alterations are seen in EEG, fMRI and DTI data. We suspect that our misclassified “healthy” may be one of these individuals who may have suffered one or more concussions previously and are exhibiting persistent changes. We did not request our subjects’ concussion history since there is considerable evidence showing that self-reported histories are prone to errors [70, 71]. Ultimately, the resolution of this puzzle requires growing our relatively shallow dataset with a larger sample that encompasses the full range of diversity present during the acute post concussion phase.

Another fascinating question emerging from our study is: how is *ConcNet* able to leverage raw, highly nonstationary, EEG time series data to distinguish between concussed and healthy adolescent subjects? Given that an LSTM-based network’s main advantage is that it can identify temporal correlations and causal relations within a sequential data stream, we suspect that *ConcNet* may be picking up alterations in the temporal dynamics of brain due to mTBI. There is evidence showing that concussions do just this: By applying the sliding window analysis to resting state fMRI data, Muller and Virji-Babul [79] identified three distinct brain states and showed that while healthy adolescents switched dynamically between the three brain states and spend approximately the same amount of time in each brain state, concussed adolescents spent the majority of time in only one brain state. The possibility of using neural networks, like our LSTM-based *ConcNet*, to probe brain dynamics is an exciting new frontier that we are starting to explore.

To summarize, in this paper, we introduced *ConcNet*, a high-performing deep learning Long Short Term Memory (LSTM)-based recurrent neural network that is able to distinguish between healthy and acute post-concussed adolescent athletes using only a short (i.e. 90 seconds long) sample of resting state EEG data as input. The athletes were neither required to perform a specific task nor subjected to a stimulus during data collection, and the acquired EEG data was neither filtered, cleaned of artefacts, nor subjected to explicit feature extraction. The classifier’s final network architecture is shown in Figure 3. *ConcNet* was trained and tested using a total of 216 EEG datasets, obtained from 27 male, adolescent athletes with sports related concussion within 1 month of injury, and 280 datasets from 35 age and sex matched healthy athletes. Using a stratified balanced sampling strategy, we trained an ensemble of 100 independent networks in order to carry out a detailed assessment of the classifier. *ConcNet* achieved median values of 0.917, 0.900 and 1 on *Accuracy, Specificity* and *Recall* respectively (see Table 4 for more details). The magnitudes of all three are reassuringly high. These metrics do, however, depend on the adopted threshold for classification. Presently, we use a threshold of 0.5 to distinguish between concussed and healthy, with participants who are assigned a probability score greater than the threshold (i.e. *P*_mTBI_ > 0.5) being classified as concussed. Adopting a different threshold, like *p*_mTBI_ > 0.4, increases the metric values and reduces the associated scatter. We also evaluated *ConcNet*’s Area Under the Curve, a performance metric that does not depend on the adopted threshold for classification. The resulting ensemble-median value of 0.971 confirms that in the context of the population group that *ConcNet* was trained to classify, the classifier does very well at distinguishing between the concussed and the healthy subjects. This is the first instance of a high-performing, objective, brain-based, platform for automatic classification of concussion at an individual level that relies only on easy to acquire resting state EEG data. It represents a key step towards addressing one of the fundamental challenges in concussion management: the dependence on highly subjective self-reporting of symptoms when determining whether an athlete has suffered a concussion or has recovered from one.

## 4 Methods

### Participants

The EEG recordings used in this study were collected from a total of 62 male adolescent athletes. Of these, 27 individuals [mean age = 13.5 years; standard deviation (SD):± 2.6 years], had suffered a sport-related concussion, were within one month of injury, and met the concussion diagnostic criteria consistent with the Berlin consensus statement [66]. The other 35 were healthy, active, age- and sex-matched cohort of athletes [mean age: 14.7 years; SD:): ± 2.1 years]. We chose the time window of one month based on the 2017 Berlin Consensus Statement’s expected time for clinical recovery after injury in children and adolescents [80]. Here, concussion is defined as a traumatic brain injury caused either by a direct blow to the head, face, neck, or elsewhere on the body that leads to an impulsive force transmitted to the head, resulting in changes in one or more of the following clinical domains: (i) physiological (e.g. neck pain, balance problems, headache, fatigue), (ii) cognitive (e.g. difficulty with attention, feeling in a “fog”), (iii) emotional (e.g. irritability, sadness) and (iv) behavioural (i.e. sleep/wake disturbances). In the case of the concussed participants, the date and time of the direct blow was documented by the team coach as per the consensus statement, and the team physician or a physician with expertise in concussion made the diagnosis of concussion again as per the Berlin consensus statement. Individuals with focal neurologic deficits, pathology and/or those on prescription medications for neurological or psychiatric conditions were excluded from this study. Our study was approved by the University of British Columbia Clinical Research Ethics Board (Approval number: H17-02973). All participants provided assent and the adolescents’ parents gave written informed consent for their children’s participation under the approval of the ethics committee of the University of British Columbia and in accordance with the Helsinki declaration.

### EEG recordings

#### Data collection

Five minutes of resting state EEG data were collected while participants had their eyes closed. We used a 64-channel HydroCel Geodesic Sensor Net (EGI, Eugene, OR) connected to a Net Amps 300 amplifier [81]. The signals were referenced to the vertex (Cz) and recorded at a sampling rate of 250 Hz. The scalp electrode impedance values were typically less than 50 *k*Ω. To eliminate any transients at the start and the end of a data collection session (due, for example, to subjects taking a few seconds to settle down, etc.), 1000 data points were removed from the beginning and the end of each time series; this corresponds to removing data with a total duration of 8 seconds. We did not filter or otherwise clean the data to remove artefacts due to line noise, eye blinks and motion as is typically done (c.f. Porter et al. [82], Rotem-Kohavi et al. [83, 84], Munia et al. [19]).

#### Data segmentation and augmentation

We segmented the 64-channel EEG data into three 90 seconds long consecutive sets of raw90-rsEEG. We further augmented our data compilation by extracting, from each subject’s five minutes long 64-channel data stream, five additional 90 seconds long, synchronous segments with randomly chosen starting points. Thus, in total, we extracted 8 sets of raw90-rsEEG recordings per participant. EEG signals are well known to be strongly non-stationary on timescales greater than ∼0.25 s, exhibiting significant variations, and even shifts, in the statistical properties of the signal over time [61, 62]. This behaviour poses a significant challenge for extracting stable features from the signal as well as for designing reliable brain-computer interface systems within the context of real environments [85–88]. In the present study, we leverage this behaviour to generate multiple, effectively independent, realizations of the raw90-rsEEG datasets from each of the participants.

Our choice to use 90-second long segments was the result of a compromise between wanting to use a short EEG sample on one hand, and ensuring that the sample is long enough to capture very low frequency brain activity, i.e. oscillations with frequencies down to ∼ 0.01 − 0.02 Hz (also known as infraslow oscillations or ISO). The latter was motivated by two considerations: (a) EEG studies find that, compared to healthy controls, concussed athletes exhibit altered activity in the delta band (0.5 − 4 Hz) [19, 33, 89], with increasing divergences towards to lowest frequencies. (b) Among adolescents, one of the resting state networks strongly impacted by concussion is the default mode network (DMN) [20, 48]. This network, as well as several other resting state networks, support very low frequency, large amplitude oscillations [90]. The power in these frequency modes has been shown to scale approximately inversely with frequency [91]; this 1*/ f* background has a profound impact on neuron dynamics and communications [92].

### Behavioural Data

When the EEG data were collected, the number of symptoms and symptom severity of each of the concussed subjects were also evaluated using either the Child Sports Concussion Assessment Tool 3 (Child SCAT3), if the injured athlete was younger than 13 years of age (https://bjsm.bmj.com/content/bjsports/47/5/263.full.pdf), or using the Sports Concussion Assessment Tool 3 (SCAT3), if 13 years or older (https://bjsm.bmj.com/content/bjsports/47/5/259.full.pdf). Both SCAT3 and Child SCAT3 are standardized concussion and concussion symptom assessment tools. SCAT3 lists 22 symptoms that can be scored from 0 (none) to 6 (severe) while Child SCAT3 lists 20 symptoms with scores ranging from 0 (none) to 3 (often). The overall symptom severity score of the injured athletes is the sum of the individual symptom ratings. SCAT3 allows for a maximum score of 132 while the highest possible Child SCAT3 score is 60.

### Network Architecture

As noted in §2, we had previously experimented with several popular machine learning classification algorithms, including the Support Vector Machine (SVM) algorithm [41] and the simplest type of multilayer perceptron (MLP) neural network [45]. The latter was a three-layer network consisting of an input layer that receives the data, a single hidden layer that processes the data, and an output layer that classifies based on the results from the hidden layer. The input for both attempts consisted of a set of statistical measures (hereafter, *features*) extracted from each individual’s set of 64 resting state EEG time series data. Neither of these two attempts yielded classification accuracy above 65%, a finding that was not a complete surprise. It is well known that measures extracted from different segments of time duration Δ*t*, extracted from even a single non-stationary EEG time series collected at one sitting, can vary significantly [61, 62, 87]. However, prior to extracting the features, we had filtered the data [82–84] to remove signal drift, an approach that is commonly adopted to mitigate weak non-stationary behaviour. This works well enough when one is interested in *population-level* differences between concussed and healthy groups [24, 28, 47, 48]. However, the distribution functions of features for the healthy and concussed individuals overlap, which hampers attempts to use the features to assess *individual* subjects. The problem is further compounded by the fact that EEG time series is strongly non-stationary and simple procedures, like filtering, do not completely remove the non-stationarity of features. This tendency is the source of the difficulties faced by researchers pursuing reliable brain-computer interfaces [86, 88]. In light of this, and in the interest of ensuring that the data processing pipeline is as simple as possible, we felt it best to use the observed (raw) EEG signals and rely on a deep network to learn the signals.

As for the choice of the deep learning network architecture, we took our cue from the brain-computer interface (BCI) literature and adopted a recurrent neural network architecture comprising two layers of bi-directional Long Short Term Memory (LSTM) [63, 64] units. In the context of EEG-based BCIs, Recurrent Neural Networks (RNN) — and in particular, a 2-layer bi-directional LSTM network — have been found to be better at accommodating non-stationary behaviour of EEG time series and consequently, yield higher accuracy in classification tasks [88]. The key feature is the inclusion of *memory units*. As a sequence is processed, the memory of the results associated with prior time points persists in these units and informs the processing of current and future inputs. This facility for capturing long-term dependencies is the reason why LSTMs are better able to learn time-series data due to complex nonlinear dynamics. Our network employs bi-directional memory units [93]. These integrate information from both past and future time steps to compute the activation signals as well as the corrections that are applied to the weights during the back propagation step. Bi-directionality enhances network stability, speeds up training, and results in improved performance in comparision to uni-directional networks [94]. Stability is especially important when training with sparse datasets, as in our case. This coupled with the fact that an LSTM network is better able to handle sequential data made this architecture an obvious first choice.

Apart from the LSTM layers, our network also incorporates fully connected (FC) layers. Each node on a FC layer receives input from all of the nodes on the preceding layer. These inputs are multiplied by the weights and then summed along with a constant term. The result is then mapped to an output using a activation function. We have opted to use the *ReLU* (Rectified Linear Unit) activation function in our network because it has a much greater dynamic range than offered by other commonly used activation functions and therefore enables efficient learning in deep learning environments, and because its non-linear character facilitates the learning of complex dependencies within the data.

### Network Training and Validation

#### Stage 1

For the *blinded, one-shot* protocol used in this stage (see §2), *ConcNet* was trained using only a subset of the available dataset: that is, all raw90-rsEEG segments harvested from 44 (25 healthy and 19 concussed) of the total of 62 subjects; the remaining were reserved for the blind test. We divided this training data into two groups, the training and the validation sets, using a modified 80:20 prescription, with the training set comprising all raw90-rsEEG segments from 15 randomly chosen concussed subjects and an equal number of randomly chosen healthy subjects. The balance of the data were used for validation. We did not allow for any overlap between the two sets; all the raw90-rsEEG segments from any one subject either belonged to the training or the validation set, but not both. Since the data segmentation described in §4 provided eight data segments per subject, the training:validation split resulted in 120 segments each from concussed and control subjects for training, and 32 concussed and 80 control segments for validation.

To train the network, we adopted the *mini-batch* approach: The training proceeded in steps, each using 20 randomly selected, raw90-rsEEG segments, which were processed through *ConcNet*. The difference between the labeled classification and that predicted by *ConcNet*, summed over the 20 samples in the mini-batch was used to define the *loss function*; the smaller the loss, the better the performance. During the subsequent *backpropagation* step, all the weights in the network were updated proportional to the gradient of the loss function with respect to each weight. Validation of the training process was done after every five mini-batches, providing a metric to assess the progress of the training. These steps were continued till all the training samples had been exhausted. This marked the end of an *epoch*. At this point, the training data was reshuffled (i.e., another set of 15 concussed and 15 healthy subjects were drawn from the available data) and the mini-batch training process was repeated. The cycles were repeated for 25 epochs, by which time the loss function had levelled off, and further training resulted in little or no improvement of performance. At this point, the loss curves from training and validation were used to tune the hyperparameters. For *ConcNet*, these hyperparameters included the number of LSTM layers, the number of LSTM units per layer, the number of fully connected layers and the number of nodes in each, the regularization scheme and dropout rate used, the choice of optimizer, and the learning rate. For *ConcNet*, we adopted *Adam*, an adaptive learning rate optimization algorithm introduced by Kingma & Ba [95], which has been shown to be matched for non-stationary objectives, and works well even with noisy and sparse datasets.

We repeated the process described, including the continued tuning of the hyperparameters via an iterative search, while monitoring both the training and the validation loss curves. Training was stopped when the loss curves started to decrease monotonically and crossed a preset threshold (< 10^−4^ in our case). We did not pursue further explorations of nearly degenerate network configurations.

The full set of training iterations was carried out on a single GPU assisted cluster hosted by *Compute Canada* and took less than 30 minutes. An attempt to train the network on desktop computer equipped with a single GPU took about the same time. Given the file size and computation requirements of the training and validation datasets, the training stage needs at least one GPU, either in a compute cluster or even in a desktop computer. However, since the test data sample size is much smaller, testing the trained network could be done even on a laptop without GPU assistance.

With the hyperparameters satisfactorily tuned, the present Stage 1 *ConcNet* was tested, as noted in Section 2, via *“blinded”, one-shot* protocol, where the test data were provided to the software team without labels. The test segments were sourced from subjects that *ConcNet* had never been exposed to previously nor was it taught the data after the fact. The network results were recorded, and then the labels were unmasked for analysis.

#### Stage 2

During this stage, our aim was to optimise and simplify the network architecture as much as possible while maintaining performance. All of the data was made available and used to construct three groups: a training data set comprising the eight raw90-rsEEG segments from each of 21 randomly selected concussed and 21 randomly healthy participants, for a total of 336 samples; a validation set consisting of the eight raw90-rsEEG from each of the 3 concussed and 3 healthy participants, again randomly selected from the balance of the participants (48 samples in total); and a testing set made up of the rest. This is roughly an 80:10:10 division within the context of a stratified balanced sampling strategy, a commonly used approach for mitigating potential biases that may arise when the number of members in the groups under consideration are dissimilar. In the present instance, we require that the number of concussed and healthy participants in the training and validation sets are matched. The training and validation procedure was similar to that of Stage 1. Once training was completed, the performance was assessed using the test dataset. When faced with degenerate network configurations, we preferred those with fewer layers.

#### Stage 3

To further assess *ConcNet*’s performance as well as quantify the uncertainties in the performance metrics, we trained 100 independent networks with identical architecture and hyperparameters to final *ConcNet* configuration established at the end of Stage 2. Each network was trained using the same procedure adopted for Stage 2. Each of the 100 networks was trained using a separate, randomly selected realization of training, validation and testing sets. For this reason, the internal weights of each of the networks at the end of training, and hence their individual performance, are expected to vary from network to network, thus yielding the uncertainties associated with each of the performance metrics. Over the course of Stage 3, we maintained a detailed log of which concussed and healthy participants were being used to train, validate and test each network, as well as the resulting classification scores and the predicted labels. Here, the classification scores refer to the probabilities output from the final softmax layer. We also calculated the performance metrics for each of the networks. This type of assessment is critical for determining how well *ConcNet* would generalize to a larger, more heterogeneous datasets.

## Acknowledgements

The authors would like to thank Nazim Madhavji (Computer Science, Western University) and Kwang Moo Yi (Computer Science, University of Victoria) for valuable advice and insightful discussions. We would also like to acknowledge all the athletes from Seafair Minor Hockey Association and Richmond Football Club who participated in this study. This research was enabled in part by support provided by WestGrid (www.westgrid.ca) and Compute Canada (www.computecanada.ca).

## Author contributions statement

A.B. and D.T.H. conceived the work; N.V.B. and L.B. collected and prepared the data; A.B., K.T., B.F., M.B. and A.G contributed to the design of the network; K.T., B.F., M.B. and A.G. trained and tested the network; A.B., N.V.B., K.T., and D.T.H. interpreted the results; A.B. and K.T. drafted the paper; A.B., K.T., D.T.H. and N.V.B. substantially revised the paper. All authors reviewed the manuscript.

## Additional information

### Competing interests

The authors declare no competing interests.

## Notes

### Competing Interest Statement

The authors have declared no competing interest.

